# ntsm: an alignment-free, ultra low coverage, sequencing technology agnostic, intraspecies sample comparison tool for sample swap detection

**DOI:** 10.1101/2023.11.01.565041

**Authors:** Justin Chu, Jiazhen Rong, Xiaowen Feng, Heng Li

## Abstract

**Background:** Due to human error, sample swapping in large cohort studies with heterogeneous data types (e.g. mix of Oxford Nanopore, Pacific Bioscience, Illumina data, *etc*.) remains a common issue plaguing large-scale studies. At present, all sample swapping detection methods require costly and unnecessary (e.g. if data is only used for genome assembly) alignment, positional sorting, and indexing of the data in order to compare similarly. As studies include more samples and new sequencing data types, robust quality control tools will become increasingly important.

**Findings:** The similarity between samples can be determined using indexed *k*-mer sequence variants. To increase statistical power, we use coverage information on variant sites, calculating similarity using a likelihood ratio-based test. Per sample error rate, and coverage bias (*i*.*e*. missing sites) can also be estimated with this information, which can be used to determine if a spatially indexed PCA-based pre-screening method can be used, which can greatly speed up analysis by preventing exhaustive all-to-all comparisons.

**Conclusions:** Because this tool processes raw data, is faster than alignment, and can be used on very low coverage data, it can save an immense degree of computational resources in standard QC pipelines. It is robust enough to be used on different sequencing data types, important in studies that leverage the strengths of different sequencing technologies. In addition to its primary use case of sample-swap detection, this method provides other useful information useful in QC, such as error rate and coverage bias, as well as population-level PCA ancestry analysis visualization.

## Introduction

Large-scale sequencing studies often have robust error reduction strategies, though none are immune to human error. If sample swaps occur it can be trivial to detect known contaminants using sequence classification tools [1,2], or distance-based analysis such as MASH [3], however sample swaps in intra-species studies can be quite a bit more difficult to detect as the sensitivity would be overwhelmed by the high degree of similarity between the samples unable to distinguish between differences cause by sequencing errors or other artifacts like batch effects.

For same-species sample swap detection, using prior knowledge of variants with a minor allele frequency (MAF) ideally near 50% within the population can help increase the sensitivity of the analysis to only differences between individuals. For detecting sample swaps between the same species, in particular humans, multiple tools have been developed utilizing variant sites [4–11]. These methods rely on upstream alignment, sorting and indexing of the data, many initially require a variant calling pipeline as well, though at least newer methods such as Somalier [11] do not require a variant calling set working directly on alignments. In addition, even these methods may be overwhelmed when comparing low coverage or However even these methods may be overwhelmed when comparing heterogeneous data types such as Illumina sequencing [12] and Oxford Nanopore sequencing [13], or specialized library preparation methods upstream of sequencing such as Hi-C [14] or 10x Chromium [15] linked read sequencing data.

Rather than determining sample swaps after the alignment, sorting and indexing of the sequence data, it may be ideal to detect sample swap or other issues at the furthest possible upstream analysis point as to minimize extraneous computational costs. It may be argued that alignment may not incur any additional analysis costs as such things may be part of the downstream analysis anyway, however, studies that do not require alignments exist. For example, the Vertebrate Genome Project (VGP) [16] assembles PacBio High-Fidelity (HiFi) reads and Hi-C short reads in the lack of a known reference genome; the Human Pangenome Reference Consortium (HPRC) [17] does assembly without aligning them to the reference human genome. In addition, data specialized for other purposes than variant calling is also difficult to use in these pipelines. In light of these issues, we seek to create a tool that generically detects sample swaps and that are convenient to use upstream of any analyses.

We have created a tool for fast sample swap detection on raw whole genome sequencing data, agnostic of sequencing technology. As it uses only *k*-mer counts in the analysis and lacks the requirement of any alignment and sorting, it is unrivaled in speed compared to traditional alignment based methods, and can function on any kind of sequence data even at very low coverage data as long as the raw data is mostly uniform in coverage. In addition, the *k*-mer count information also provides the extra quality control utility such as error rate estimation and PCA population analysis to determine sample population of origin.

## Methods

### Availability

**Project name: ntsm**

**Project home page: https://github.com/JustinChu/ntsm**

**Operating system(s): Linux**

**Programming language: C++**

**Other requirements: TBD**

**License: MIT**

### Algorithm Overview

We developed ntsm focusing on minimizing upstream processing as much as possible. It starts by counting the relevant variant *k*-mers from a sample only keeping information needed to perform the downstream analysis. The counting can be set to terminate early if sufficient read coverage is obtained. Once generated the counts can be compared in a pairwise manner using a likelihood-ratio based test. During this, sequence error rate is also estimated using the counts. The number of tests can be reduced by specifying an optional PCA rotation matrix and normalization matrix adding a prefiltering step on high quality samples. Finally, matching sample pairs are outputted in a tsv file.

### Selection of variant sites and *k*-mers for human samples

For tools of this nature to function effectively, it is important to select a robust set of polymorphic sites. In our case we attempt to select sites that primarily have 2 variants within the population, that occur at high (ideally near 50%) population frequencies. For our *k*-mer based method, the sites must also not contain repetitive *k*-mers, as the coverage influences the computed confidence of our statistical test. Finally, in our selection of site for human samples we included other criteria for selection (Figure 1) which, while helpful for various reasons, are less important for our tool to function properly.

**Figure 1.**
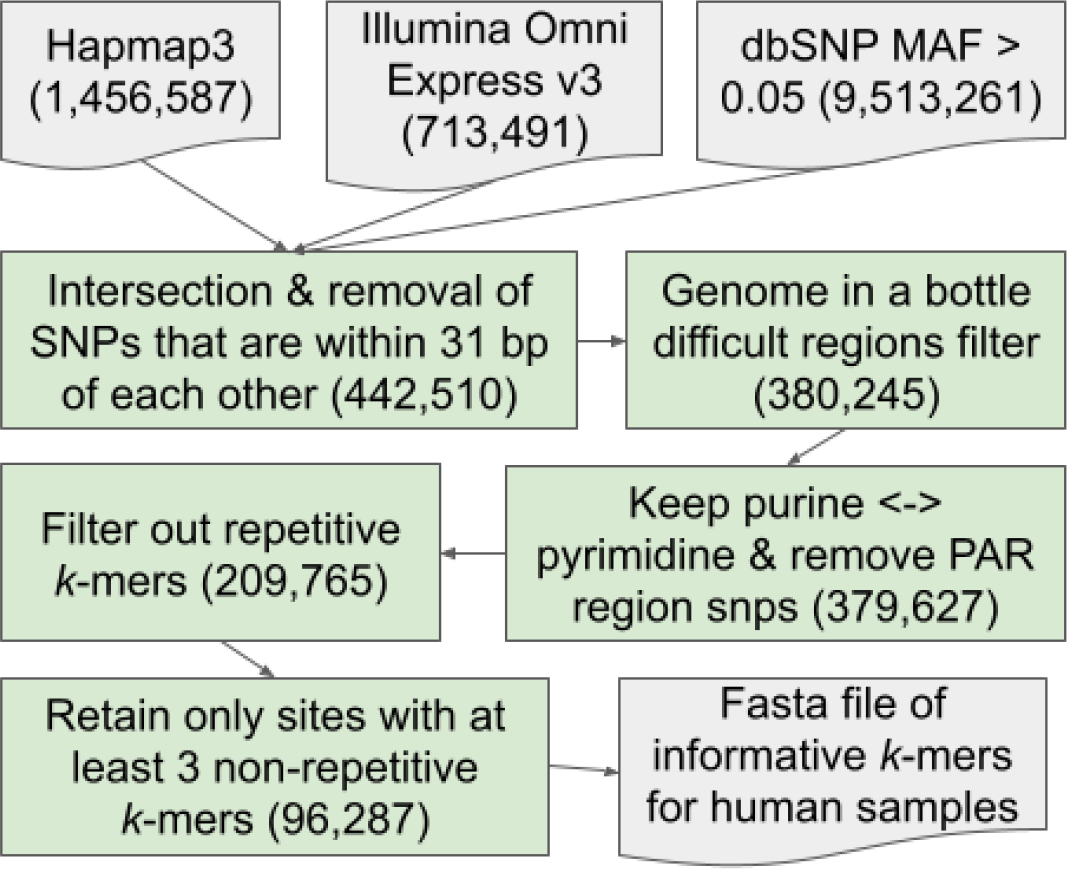
The selection method for informative human polymorphic sites used in *k*-mer counting based sample swap detection.

The polymorphic sites for human data are initially derived from an intersection of Hapmap3 [18] and Illumina Omni Express v3 [19] SNP chip sites, selected for their practical reliability allowing for possible comparisons of data using only these sites. These candidate sites are then filtered by cross-referencing the dbSNP [20] database to retain sites with a minor allele frequency (MAF) > 0.05. Any sites within 31 bp of each other are also filtered out, as we use *k*-mers that need to be mostly independent of each other in our analysis. Next, we filtered the regions by difficult regions as determined by the Genome in a Bottle Consortium [20,21]. We then keep only purine to pyrimidine (A or T to G or C) variants, as final insurance against possible human error influencing this tool.

We then process each site pulling out the 19-mers within a 31bp window for each variant and align them to hg38 using bwa aln [22] to find any 19-mers that align multiple times with at most 1 mismatch to ensure we are not using any repetitive 19-mers. Any sites with at least 3 non-repetitive 19-mers within the window are kept, resulting in a final total of 96287 sites. We expect that any similar procedure to create sites for another organism will benefit from a similar filtering step to minimize the effects of repetitive sequences. As applications for human samples are expected to be quite common, we have provided the sequences for these sites with respective identifiers (rsIDs) along with our tool.

### Generation PCA rotation matrices for human samples

In addition to variant sites sequences themselves, ntsm can optionally use population derived PCA rotational matrices which can help speed up comparisons of a large number of samples. We provide a python script that utilizes pandas [23] and scikit-learn [24] for those who wish to generate their own rotational matrices from a multiVCF file.

Using the multiVCF file from the 1000 Genome project [25], we generated a matrix of samples to our selected variant sites above (0 for homozygous A/T allele, 0.5 for heterozygous alleles, and 1 for homozygous C/G alleles). This is followed by normalizing the matrix by the standard deviation for each site and we keep the normalization vector in a file. This sample to variant matrix undergoes decomposition into principal components (Figure 2), though instead of being concerned with the principal component values themselves, we are primarily interested in keeping the rotational matrices for a number of the most significant components, and using them to project the new sequences onto.

**Figure 2.**
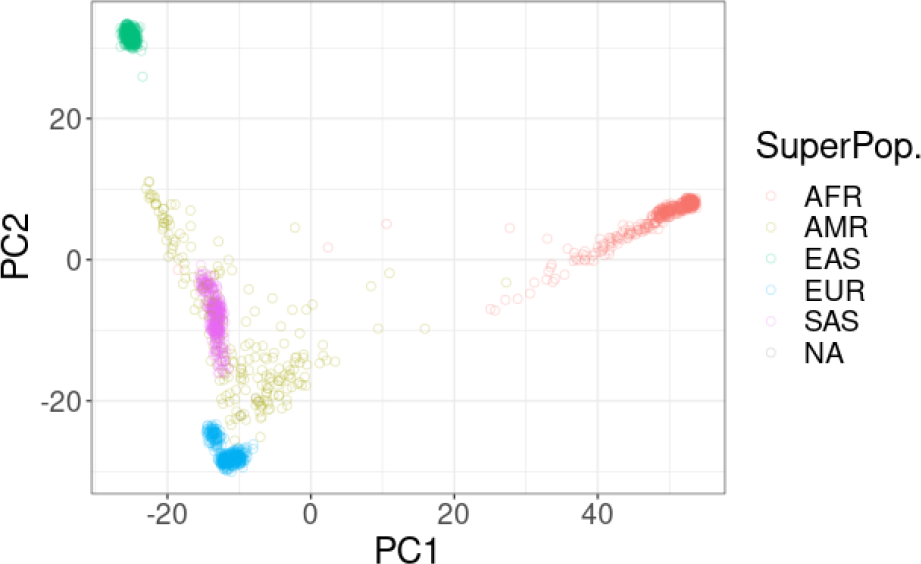
A population PCA of the first 2 components using the variant human sites selected and the 1000 Genomes Project multiVCF dataset. Each color represents a different labeled superpopulation group.

### Implementation Details

#### Variant *k*-mer Counting

Paired variant sequences (1 file for C/G allele and 1 file for A/T variants) are stored in fasta files before being loaded. These alleles are then broken into *k*-mers and hashed using an invertible hash function into a hash table [26]. A sliding *k*-mer window for each allele is used, allowing for some redundancy to compensate for sequence errors. Input sequences in fastq format are then read, broken in *k*-mers and also hashed [27], then subsequently checked for existence in the hash table. If they exist, then the occurrence count for that *k*-mer increments by one.

Optionally, the total number of *k*-mers that match a site can be used as criteria for early termination to save computational time, as only fairly low coverage is needed to perform accurate classification. For each allele, only the highest count of the sequence is outputted in the results counts file. The counts are a simple tab-separated (TSV) file with 3 columns for the site identifier, allele A/T count and allele C/G count. Multiple files and threads can be used on the same instance of the counting tool for speed, or run separately on different files and later merged.

#### Sequencing error rate estimation

Error rate can inform the user of the viability of the dataset and inform the user as to why downstream applications may be performing poorly. In our case, it may help explain why a sample swap signal is weak.

To estimate the error rate, our tool records the count of found *k*-mers *b* from our list of alleles and the number of *k*-mers *m* processed. Assuming that the data is randomly sampled from the a human genome and that least one of the allele sequences are present in the dataset, we can estimate using the number of expected *k*-mers *n* given the diploid genome size *g* and the number of distinct *k*-mers *d* in our *k*-mer set using with the following formula:

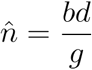

Using *n*, we can use maximum likelihood estimation [28] (MLE) to derive the estimate the expected similarity *p*:

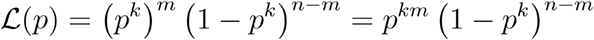

Working in log space will make our MLE derivation easier,

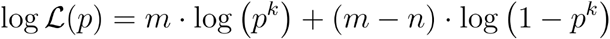

Thus,

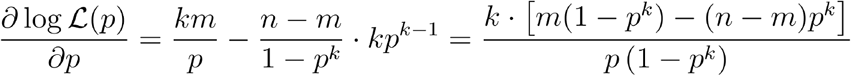

The maximum likelihood estimate of *p* is obtained when *∂* log ℒ / *∂p* = 0. Thus,

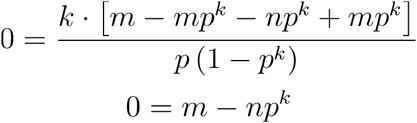

Finally, similarity is formulated as:

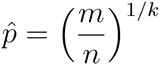

Error rate is merely the inverse of similarity,

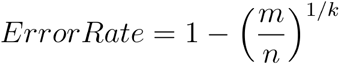

We note that this formulation largely holds true for mismatch and small indel error types, however when large indels are introduced this formulation can become less accurate depending on how one defines the ground truth alignment used to calculate the sequence error rate.

#### Similarity score for detecting sample swaps

Rather than computing genotypes for each site, our method directly uses the counts of each allele derived from k-mer counts. Our score is derived using likelihood ratio test as a basis with 2 models assuming that either the samples are independent or if they are the same sample. To minimize the effect of missing data due to low coverage, in each pairwise analysis we remove sites with missing counts for both alleles. The score is further modulated to be extra conservative, lowering the confidence of our result when coverage of the dataset is low.

To start, we need the likelihood of one sample. Suppose there are *N* sites. For a sample at site *i*, we observe *x*_*ia*_ count of allele *a*, where *a* ∈{1, 2} for the two alleles at each site *i*. Let *p*_*ia*_ be the probability of observing allele *a* at site *i*. Then the probability of the data **x** is:

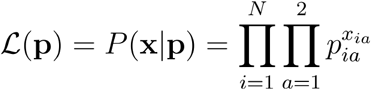

The log-likelihood is

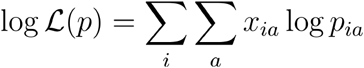

The max-likelihood (ML) estimate of *p*_*ia*_ is

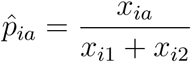

Next, for a log-likelihood ratio test, we need to compare two models:

Model 1: two samples are independent

Let *L*^*(1)*^ be the likelihood of sample 1 and so it is with sample 2. The total likelihood is *L*^*(1)*^ · *L*^*(2)*^.

Model 2: two samples are the same

In this case, we can merge all counts. Let

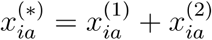

Then the probability of the two sample is

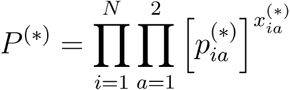

The ML estimate of 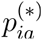 is

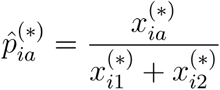

and the log-likelihood is

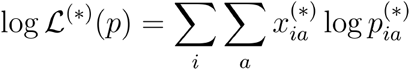

Using the two models proposed previously we can input the results into the Log-likelihood ratio test [29]. The log likelihood ratios can be used to compute a robust score metric which can determine if two samples are of the same origin. The log-likelihood formulation is as follows:

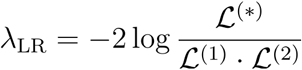

We scale by the number of non-zero sites considered *N* and the coverage *c*_*1*_ and *c*_*2*_ of both samples. The *c*_*1*_ and *c*_*2*_ are included in the formulation to reduce our confidence in the results when the coverage is low (Supplementary Figure S1), though is empirically modulated with the skew parameter *s* (default 0.2). The final score formulation is as follows:

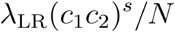

This score metric effectively accounts for lower confidence of the results when low coverage data is used.

#### PCA-based spatial-index for fast sample screening

Naively comparing samples is an all-to-all operation (*i*.*e*. O(n^2^)), in which even state-of-the-art methods such as Somalier perform when finding similar samples. Indeed, one of the key novel innovations in Somalier was the use of genome sketches to minimize the time spent on each comparison, which is admittedly extremely fast. Here we opt for a more sensitive approach that utilizes count information, which we cannot easily collapse into a sketch. This results in a notably slower single element comparison time, however the overall time complexity is still quadratic if a naive approach is performed, so any performance gains made through increasing the efficiency of pairwise operations has a limit.

If many samples are being compared, we can speed up analysis by optionally combining the concept of population level PCA analysis [30] with a spatial index data structure called a kd-tree [31], with ntsm utilizing the nanoflann implementation of kd-trees [32]. Our method of generating a population PCA is mentioned in a previous section and we provide 20 rotational matrices for the human sites. At comparison time for each sample we take the variant sites and project them onto this PCA based on an existing population structure and then use a kd-tree to index them. Using a euclidean search radius in multidimensional space, we can then select the samples that occur in the local neighborhood of the sample being tested to minimize the number of comparisons being performed. We note that in order for this method to work the data must be of high quality, missing very few sites and have very accurate allele frequencies. Thus, our implementation uses the various criteria to determine if a sample is safe to use or must undergo a large radius search or even an exhaustive search.

Our search radius in this multidimensional space is determined by 2 properties of the data - the sequence error rate (estimated via method above), and the percent of missing sites (sites with a count less than the minimum count threshold). For the first radius (default = 2), only samples with a missing site percent less than 1% and an error rate less than 1% are permitted. For the second radius (default 15) a missing site percent greater than 30% is required. Finally if the data fails all of these conditions, an exhaustive search between all pairs is performed.

#### Calculating relatedness

Our method for computing relatedness largely borrows from the exact method described in the Somalier publication [11] which compensates for loss of heterozygosity seen in many tumor samples. Our implementation uses *k*-mer counts to create rough genotyping calls and omit missing sites from the relatedness calculation. In our counts to make genotyping calls we filter *k*-mer occurrences less than 1, to compensate for *k*-mers induced by sequencing errors.

## Results

### Validation of error rate estimation

Sample swap detection works better when sequence data is largely free from errors. However, though sequencing error rate can be broadly estimated by type of technology used, a sequence based estimate of the error rate can be invaluable to troubleshoot why some samples may have stronger or weaker or associations than expected. In addition, error rate is used when screening samples on whether our PCA based index can be used and autodetection of this helps simplify the user experience.

To measure the accuracy of our error rate estimation, our ground truth was based on the alignments to the CHM13 T2T reference genome [33]. We chose this effectively haploid genome to minimize any over estimation of error due using alignment to a reference as the ground truth. We used real Illumina, Pacbio Hifi and Oxford Nanopore data for CHM13 in addition to simulated data using wgsim [34] and PBSIM [35] at differing error rates. Error rate for real data is defined after alignment and we used the gap collapsed error rate (*i*.*e*. gap collapsed sequence identity [36]) metric in this case. Gap compressed error rate does not take into account error caused by gap lengthening but still takes into account indel and mismatches.

We found that our estimates closely match the expected error rate (Fig. 3), though both the real and simulated ONT data was slightly underestimated on average, but not to a degree that makes the estimate unreliable. It is expected that error caused by indels, especially if long segments of these are present, would produce a lower calculated error rate as the formulation (see methods) for our error rate calculator assumes only mismatches can occur. That said, we expected the formulation’s the short indels to contribute to the error calculation in a way very similar to mismatches which is reflected here. We note that for our method an estimated diploid genome is needed, and for these tests the default value used was 6.2 Gb. This value of course will differ if a genome with a different size genome is used.

**Figure 3.**
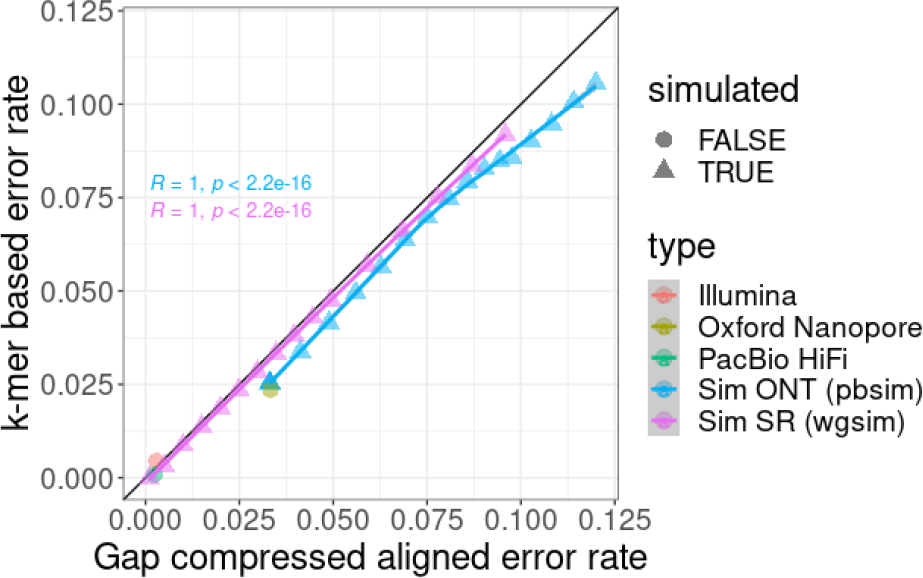
*k*-mer based error rate calculation vs Gap compressed align error rate ground truth on real and simulated CHM13 data onto the CHM13 T2T reference genome.

### PCA-based heuristic investigation

To evaluate the efficacy of our PCA-based method to reduce the number of pairwise comparisons that we perform, we use data from the HPRC (Supplementary Table S1), and run them with ntsm while providing a rotation matrix with normalization values. The properties of the data varied wildly between coverage and error rate providing comprehensive expected performance of our heuristics given different data types.

The number of missing sites greatly affects the performance of our heuristic, generally requiring a larger radius to search as the number of missing sites increase (Figure 4). We can measure the number of missing sites to apply thresholds prior to applying a search radius. The other variable determining the performance of our heuristic is the correctness of our genotype calls on our data. This is influenced by both the coverage of the data and error rate of the data. We also use our error rate estimates to determine the radius to search. The coverage is largely a function of the missing sites and found it to be a better metric overall. Because of this, high coverage data generally requires a much smaller search radius (Supp. Figure S2).

**Figure 4.**
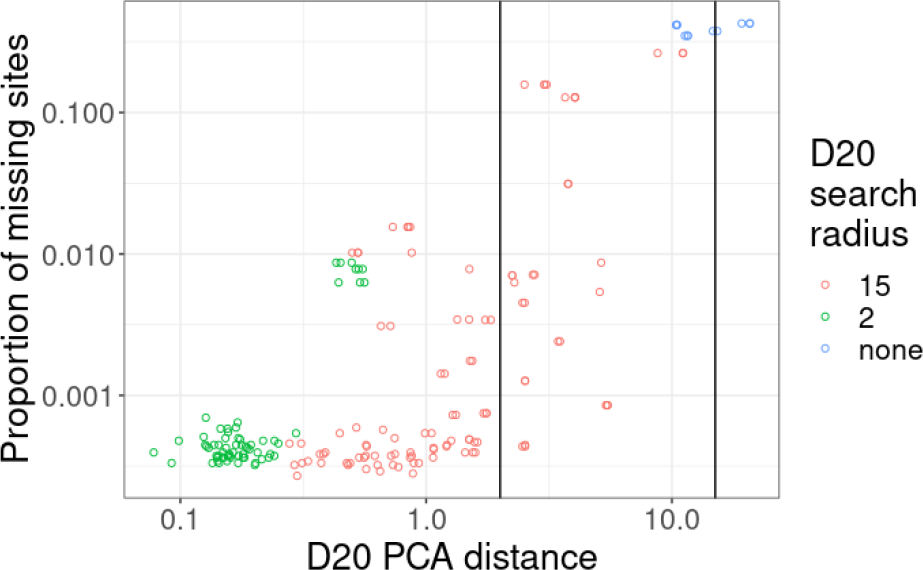
The PCA distance (top 20 PCs) between matching sample pairs vs the proportion of missing sites found in the dataset with the most missing sites. The colors (D20 search radius) correspond to the radius chosen by the algorithm for each sample pair based on the error rate and number of missing sites The vertical lines denote the radius thresholds of 2 and 15.

When the *k*-mer coverage smaller dataset of a pair of matching samples >20x, we find that >57% of the samples are set to use a radius of 2, while at a coverage >30x, we find that >93% of the samples require a search radius of 2 (the remaining datasets at search radius of 15 being higher error nanopore data). At a search radius of 15 the expected number of candidates is <25% of the total number of samples, and at 2 the expected number of candidates to search drops to <5% of possible elements on average (Figure 5).

**Figure 5.**
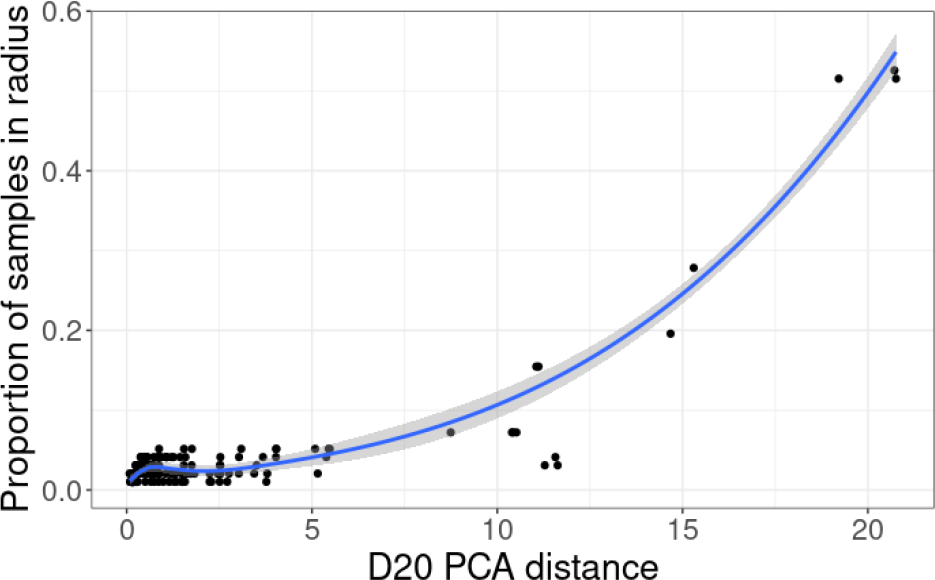
The PCA distance between each matching sample pair and the average proportion of samples of that are determined as candidates for at that radius.

### Comparisons to Somalier

To show the performance of ntsm, we sequence data from the Human Pangenome Reference Consortium (HPRC) [17] and a multiVCF file from the 1000 Genomes Project [25] featuring 3202 samples. We compare 39 samples with whole genome data from the HPRC which include sequencing data from Illumina, Pacbio HiFi, Hi-C, 10x Chromium, Strand-seq, and Oxford Nanopore platforms (Supplementary Table S1). As we expected Oxford Nanopore to be the most difficult datatype to analyze, we chose only samples that had complementary data of this type in the analysis.

#### Sensitivity and Specificity of Sample Swaps

Unlike ntsm, Somalier does not provide an automated means of determining what should be considered similar enough to consider it the same sample. However, it does provide relatedness metrics that can easily be used to threshold samples showing high degrees of relatedness and thus similarity. To determine a threshold, we ran the full coverage of each dataset for Somalier and manually picked a threshold that provided perfect separations between samples with the same origins from those with different origins (Figure 6). We determined that a relatedness value of 0.667 seemed to be a good threshold while still maintaining a high sensitivity. As mentioned in the methods section, ntsm uses a log-likelihood based score to separate samples with same origins from those with different origins, where the default value for this threshold is 0.5.

**Figure 6.**
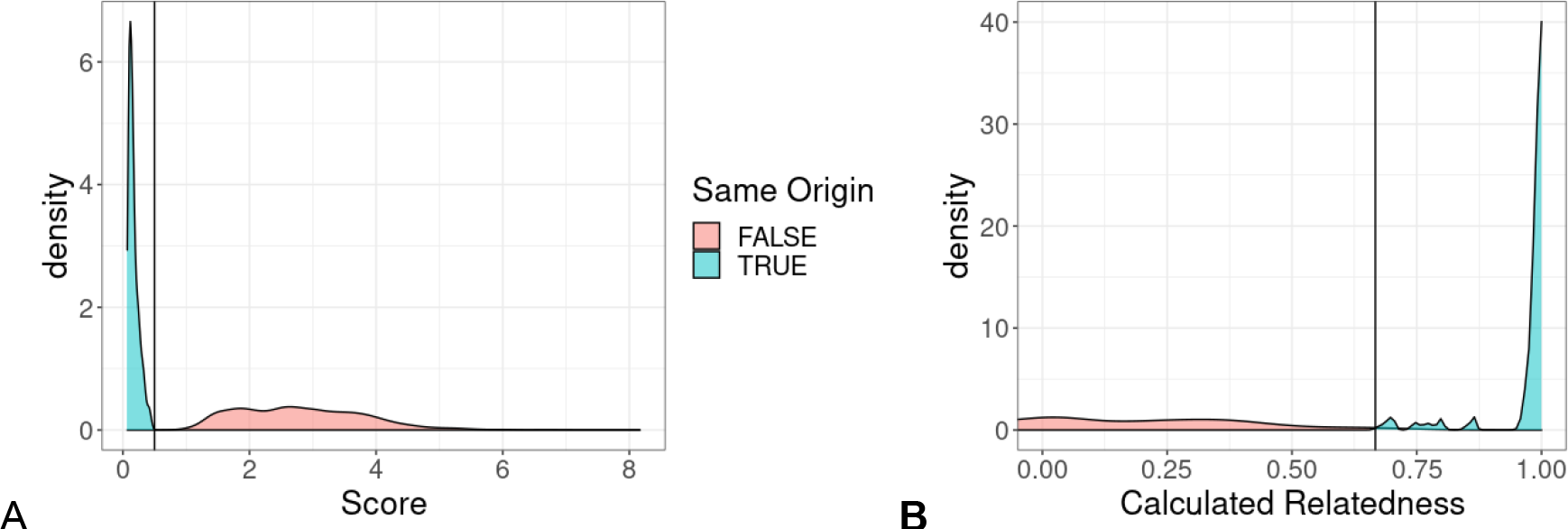
Distributions of metrics (score for ntsm (B) and relatedness for Somalier (A)) used for determining if two samples are similar. Horizontal lines are the thresholds used to determine the same sample status in later parts of the comparison (Relatedness > 0.67 for Somalier and Score < 0.5 for ntsm, respectively).

Next we randomly subsampled each dataset at different fold coverages (from 0.5x to 20x) and proceeded to see how each tool performed on this lower coverage data. We found that both tools are capable of detecting whether or samples have the same sample of origin at coverages higher than 5x, however Somalier struggled when attempting to match samples at coverage lower than 5x, producing a much higher number of false positive and negative pairs in the output (Figure 7). At sub-1x coverage, even ntsm struggled with detecting samples with the same origin, though not to the degree that Somalier struggled. In general, ntsm benefits from higher coverage as well, generating higher scores (i.e. more confident results) for unmatched samples (Supplementary Figure S1.).

**Figure 7.**
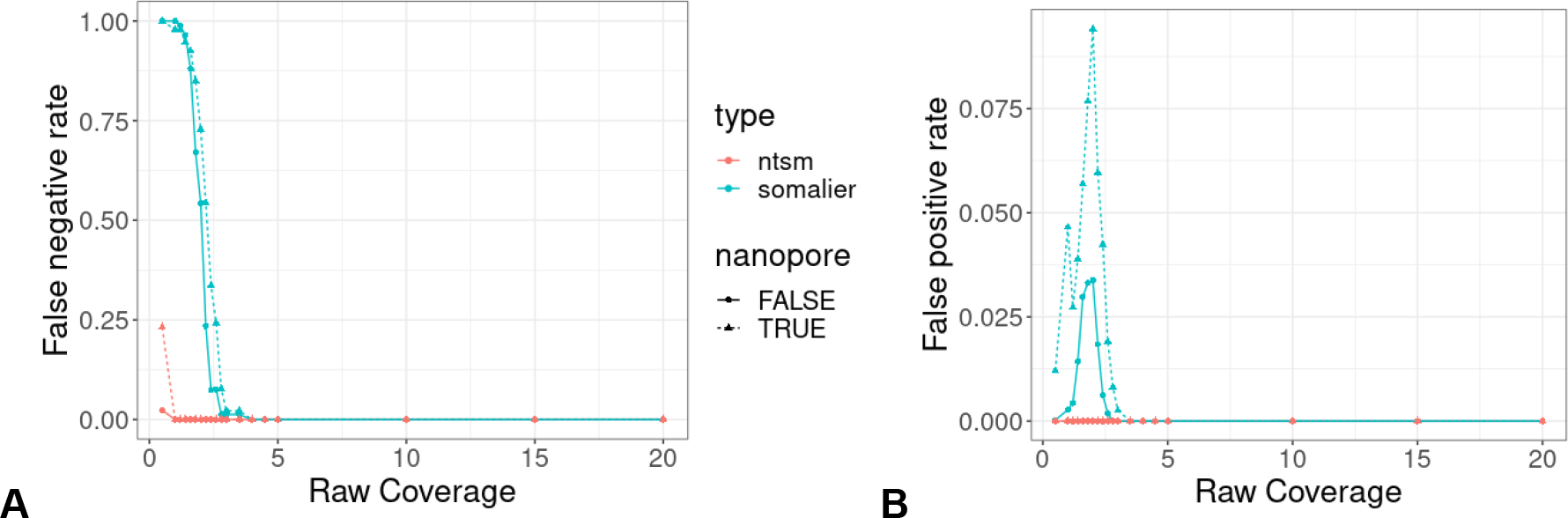
Effects of dataset coverage on classification performance on Somalier and ntsm. A. The false negative rate (FNR) of Somalier and ntsm at varying raw dataset coverage. B. The false positive rate (FPR) of Somalier and ntsm at varying raw dataset coverage. Nanopore combined with other datasets is shown separately due to the high error rate of the former.

#### Memory and time comparisons

##### *k*-mer counting (ntsm) vs alignment (somalier)

One of the primary benefits of ntsm is bypassing the alignment requirement that other tools require. However, the alternative we must perform is *k*-mer counting, which, though is fairly resource frugal, is not free. To determine the relative resource cost counting takes in comparison to alignment, we took equal coverage subsamples (2x) of our datasets (Supplementary Table S1) and ran them with ntsm and each alignment tool we used to generate alignments needed for Somalier.

We then measured the time and memory needed for each tool used (Figure 8). We found that ntsm ran at an average of ∼8 minutes, orders of magnitude less than bwa mem and minimap2 at ∼1.9 hours and 5.9 hours respectively. Memory usage is low because we are only counting a very small specific subset of *k*-mers. Note that we did not include sorting or indexing time in this analysis as we hoped to illustrate that even without this in our comparison *k*-mer counting was still much less resource intensive. Also sorting can partially be run in parallel with alignments as reads are streamed.

**Figure 8.**
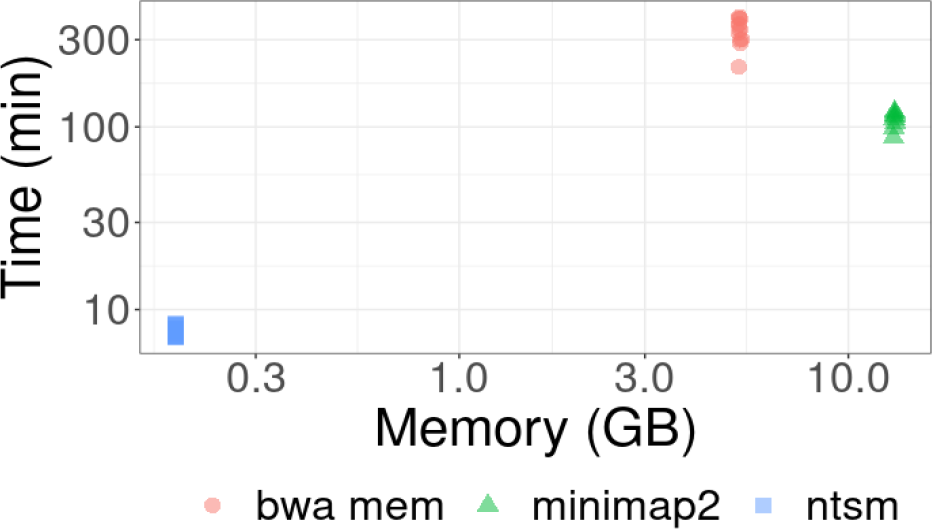
Peak memory and wallclock time comparisons of *k*-mer counting vs alignment using single thread

##### Sample comparison process

As mentioned in the method section, comparison of all samples with each other is naively an all-to-all operation and thus a quadratically scaling operation. For most studies this may still be a trivial concern but as larger and larger studies are considered, this can become an increasingly expensive consideration. To observe the computational expense of this process we took subsamples of the 1000 Genomes VCF files and measured time and memory used to compute relatedness between the samples.

As expected we observed that pairwise comparison time scaled quadratically (Figure 9), even in Somalier, albeit with each individual comparison being orders of magnitude faster than our method. Somalier, utilizing a bitvector based comparison method is much more optimized for speed than our count based method, however we show that our PCA-based screening method is still competitive. We observe here that our PCA-based screen approach actually may scale less than quadratically though it requires high coverage and low error rate data to perform this way reliably. Memory usage is largely linear relative to the data (Supp. Figure S3) as expected and there is no additional memory overhead using our PCA-based method.

**Figure 9.**
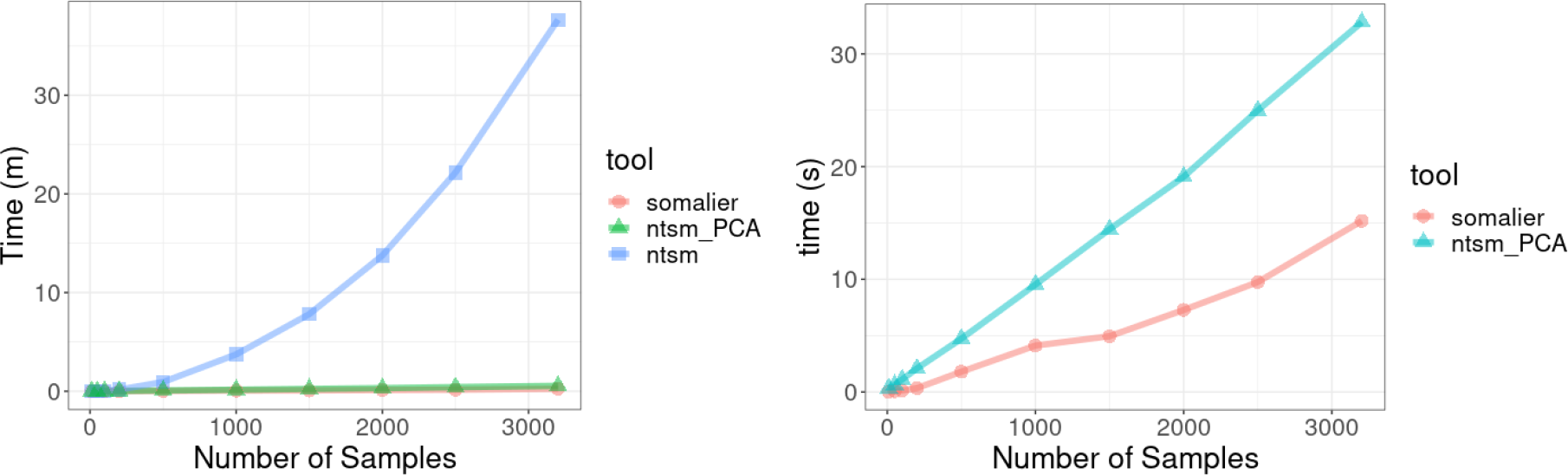
Wall clock time comparisons of somalier vs ntsm with and without PCA-based screening. A multiVCF file for 3202 samples from the 1000 Genome project was used. Somalier was capable of only using 1 thread while ntsm used 16 threads.

##### *k*-mer based relatedness calculation

To test our relatedness estimation methods we took Pacbio HiFi datasets of the parent (HG003, HG004) child (HG002) trio. Both ntsm and Somalier correctly computed the relatedness we expected; that is, parents remained unrelated (0%), while child samples showed 50% relatedness to its parents and with 100% relatedness to a technical replicate to itself (Figure 10) Though the similarity values largely agree between ntsm and Somalier, there are minor differences between our calculations. These differences likely primarily stem from the fact that Somailer and ntsm use different variant sites and that we use *k*-mer counts to create genotyping calls and omit missing sites from the relatedness calculation. The differences in our tool are likely due to the number sites used by default by somalier (total of 17,766 sites), and the sites we selected (see methods, total of 96,287 sites). In addition, some of the differences in our relatedness calculation are a consequence of omitting missing sites, which Somalier is unable to perform as it computes similarity using bit vectors of a uniform length.

**Figure 10.**
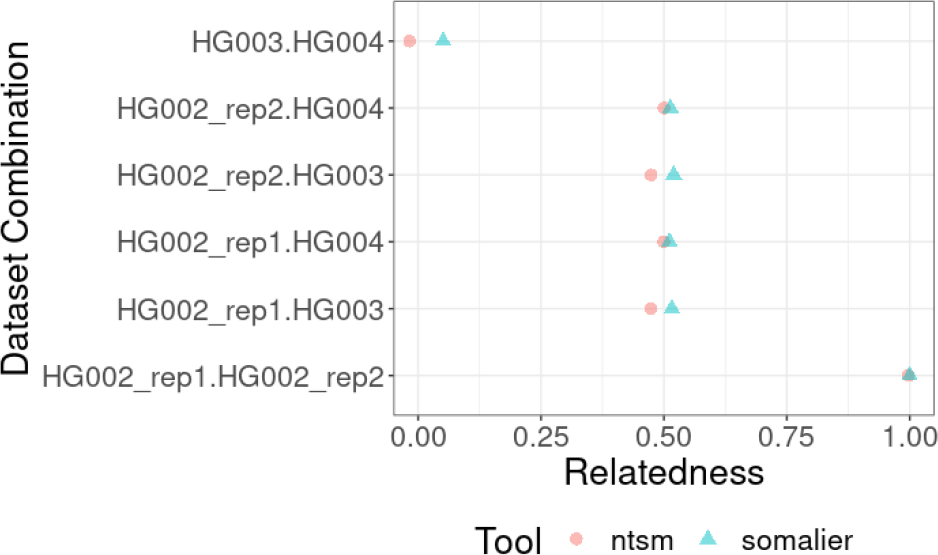
Comparison of Somalier and ntsm computed relatedness of parent child trio (HG002, HG003, HG004) from Pacbio HiFi sequencing reads. Two independent sequencing runs of HG002 are used here.

To gain a more comprehensive view on the accuracy of our relatedness calculations we checked the quality of our relatedness estimates using samples shown to be related in 1000 Genomes cohorts in addition supplemented with samples with at least 20x coverage sequence coverage of various sequencing data types (Supp. Table 1). We found that overall ntsm produces a relatedness metric closer and more tightly grouped to the expected value based on the pedigree (Figure 11). We note however this trend does not hold true for relatedness estimates involving a nanopore dataset, showing that ntsm calculates relatedness conservatively when it comes to data originating from the same sample.

**Figure 11.**
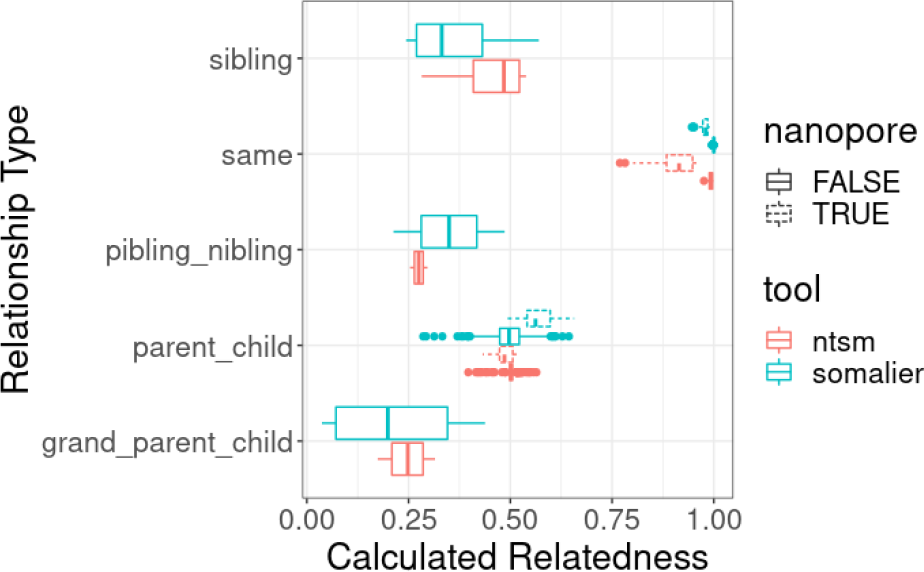
Comparison of known related samples in 1000 Genomes cohort, supplemented with samples with ∼20x coverage sequence coverage of various sequencing data types (Supp. Table 1). Relatedness calculations involving a Nanopore dataset are shown separately.

## Discussion

Here we have described NTSM, a tool designed for sample swap detection in QC contexts. The major benefits are that it uses resource frugal counts of specific *k*-mers rather than alignments decreasing overall computational costs and its capability to robustly function independent of sequencing technology type with high sensitivity on low coverage data. For large scale multi sample comparisons, we utilize a novel PCA-based spatial index heuristic screening method that greatly reduces the computational cost of comparing samples by reducing the number of candidates to compare. Overall over previous alignment based methods, we believe that ntsm could be an effective upstream tool in large scale studies, enabling robust QC and reducing the chances of error as studies become larger and incorporate more diverse sequencing data types.

### Validation and Comparison with Somalier

We compared ntsm to Somalier [4], another state-of-the-art tool designed in part for sample swap detection, using data from the HPRC [17] as well as 1000 Genome project [25]. In particular we showed that on low coverage, and high error rate data, ntsm outperforms it in terms of sensitivity and specificity, able to find 97 out of 97 matching sample pairs with no false positives at 1x coverage. In addition, though not our goal, we found that ntsm also outperformed Somalier in estimating relatedness, producing relatedness estimates closer to the known pedigrees. Somalier utilizes alignments, and we showed that alignment operations were an order of magnitude slower than our count based algorithm. However, Somalier outperformed ntsm in speed when it came to pairwise comparisons post alignment, as they used faster bit vector based comparison. PCA-based spatial index heuristic helped reduce the time complexity of our method and saved orders of magnitude by reducing the comparison, but is by is limited by the quality of the data. Overall if the goal is sample swap detection then ntsm would likely be the best choice, but if the goal is computing relatedness between all samples in a large cohort, then Somalier would be the superior choice due the speed of its pairwise comparisons.

### Reference-based vs reference-free

Our tool, unlike generic k-mer comparison methods like MASH [3], requires a set of variant sites to then derive the non-repetitive sets of *k*-mers. We recognize that a tool that uses raw *k*-mer spectrum information without a reference would be more desirable than a population/reference based method. However, any generic reference-free method that uses a *k*-mer spectrum analysis approach or similar would require much higher coverage, larger sequence differences within between samples, and very low error rates and noise. Here, the reference/population based information provides the information needed for the high statistical power and robustness of intra-species sample swap identification. Finally, beyond statistical considerations, it is computationally trivial to consider only a subset of sequences existing or not than it is to index and compare the spectrum of sequences between two samples.

### Current Limitations

Though robust enough to compare most samples originating from different sequencing technologies, not all possible sequencing data types have been tested and may not work with our method. Our statistical tests were formulated with the assumption that the input roughly originates from a whole genome shotgun sample. We have not yet tested data types such as whole exome data [37], RNA-seq [38] or ChIP-seq [39]. The data types largely differ by the extreme coverage differences between sites and the fact that more specialized sets of sites (*i*.*e*. transcribed regions) would likely need to be selected.

Our statistical test takes coverage into account to determine confidence in our test. Thus variable coverage in these datasets may pose a problem. To better adjust our method to be more robust on variable coverage, various processing methods may be worth trying. For instance, digital normalization [40] could be help for highly variable coverage data like RNA-seq.

Though our pairwise statistical tests can compensate for missing sequence, a selection of sites within genic regions, in particular sites within ubiquitously expressed genes [41] may give our method the best chance.

Though our method can be adapted to work on more than just human genomes, our method currently assumes sites with two alleles with similar frequency. Thus, detecting sample swaps of non-diploid genomes using our method will require adaptations to the models we use, but we are optimistic that principle behind it (*i*.*e*. the use of population level allele frequency information and sequence coverage information) could be used to detect sample swaps in those instances.

## Conclusions

As studies become larger and more complex, sample swaps in data are inevitable. This tool could become an integral part of upstream pipelines, robust enough to be agnostic of any sequencing technology or library preparation method. In addition, this will also reduce error when it comes to collaboration between labs and they will be able to easily match data originating from the same sample even if orthogonal sequencing technologies are used. Our novel PCA based spatial index heuristic opens the possibility of sub-quadratic comparison time complexity when comparing samples and shows its potential here, though in principle we believe the methodology can still be improved on especially when it comes to compensating for missing data. We believe our counting based alignment free methodology presented here has very little computational overhead and can readily be applied upstream to preexisting many data production pipelines.

## Supplement

### Read sequence dataset information

**Supplementary Table S1.**
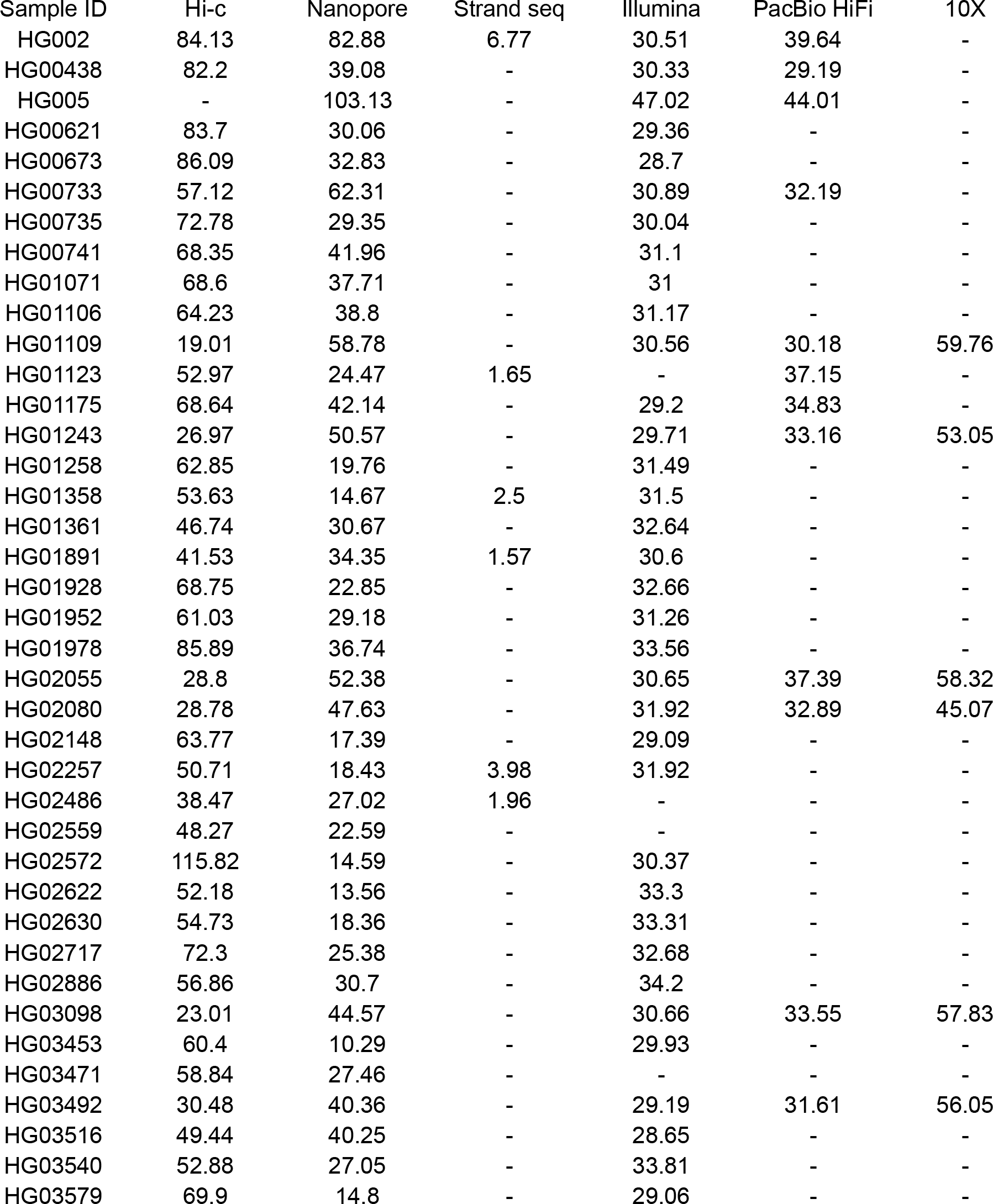
HPRC sequencing datasets used in analysis. Each column after the first represents the aligned coverage (either using minimap2 or bwa mem) of each data type available for analysis.

**Supplementary Figure S1.**
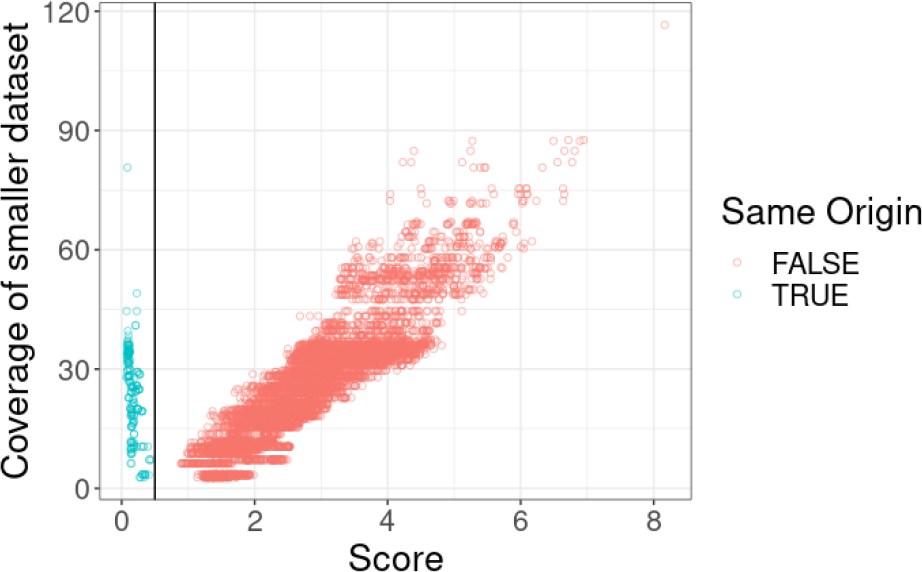
Comparison of score and *k*-mer coverage on samples from the same origin and not. Confidence in distinguishing samples increases as coverage increases, reflected in high scores the more distant they are.

**Supplementary Figure S2.**
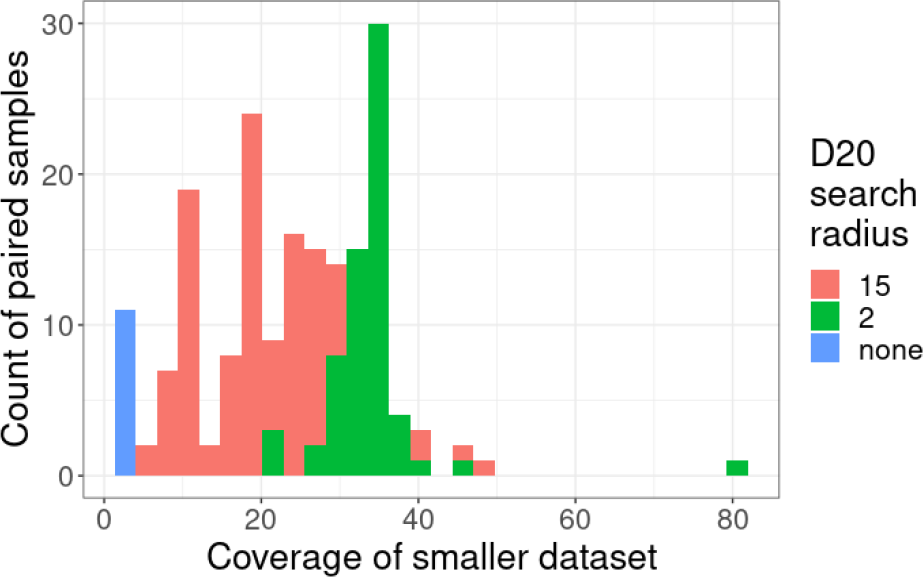
The *k*-mer coverage of the smaller sample in a matching sample pair and the number of sample pairs that use a specific search radius.

**Supplementary Figure S3.**
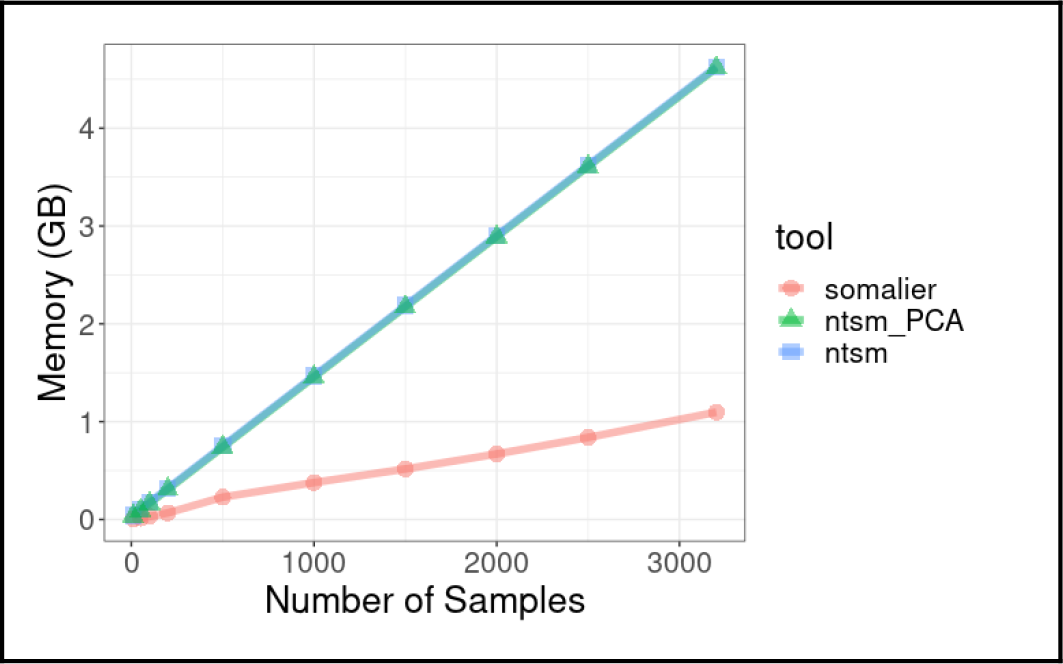
The memory of somalier vs ntsm using with and without PCA-based prescreening, processing similarity between 3202 samples from the 1000 Genome project.

## Notes

### Competing Interest Statement

The authors have declared no competing interest.

