## Supplemental Data 1 for "ntsm: an alignment-free, ultra low coverage, sequencing technology agnostic, intraspecies sample comparison tool for sample swap detection"

### Abstract

**Background:** Due to human error, sample swapping in large cohort studies with heterogeneous data types (e.g. mix of Oxford Nanopore, Pacific Bioscience, Illumina data, *etc.*) remains a common issue plaguing large-scale studies. At present, all sample swapping detection methods require costly and unnecessary (e.g. if data is only used for genome assembly) alignment, positional sorting, and indexing of the data in order to compare similarly. As studies include more samples and new sequencing data types, robust quality control tools will become increasingly important.

**Findings:** The similarity between samples can be determined using indexed *k*-mer sequence variants. To increase statistical power, we use coverage information on variant sites, calculating similarity using a likelihood ratio-based test. Per sample error rate, and coverage bias (*i.e.* missing sites) can also be estimated with this information, which can be used to determine if a spatially indexed PCA-based pre-screening method can be used, which can greatly speed up analysis by preventing exhaustive all-to-all comparisons.

**Conclusions:** Because this tool processes raw data, is faster than alignment, and can be used on very low coverage data, it can save an immense degree of computational resources in standard QC pipelines. It is robust enough to be used on different sequencing data types, important in studies that leverage the strengths of different sequencing technologies. In addition to its primary use case of sample-swap detection, this method provides other useful information useful in QC, such as error rate and coverage bias, as well as population-level PCA ancestry analysis visualization.

### Keywords

Sample mixup, Sample-swap detection, Alignment-free, Sequencing technology agnostic, Quality control, Error rate estimation, PCA-based population analysis, Low sequence coverage analysis

### Introduction

Large-scale sequencing studies often have robust error reduction strategies, though none are immune to human error. If sample swaps occur it can be trivial to detect known contaminants using sequence classification tools [1,2], or distance-based analysis such as MASH [3], however sample swaps in intra-species studies can be difficult to detect as the high degree of similarity due to being the same species can overwhelm the signal to distinguish unrelated samples, which can be further confounded by sequencing error or other artifacts caused by batch effects.

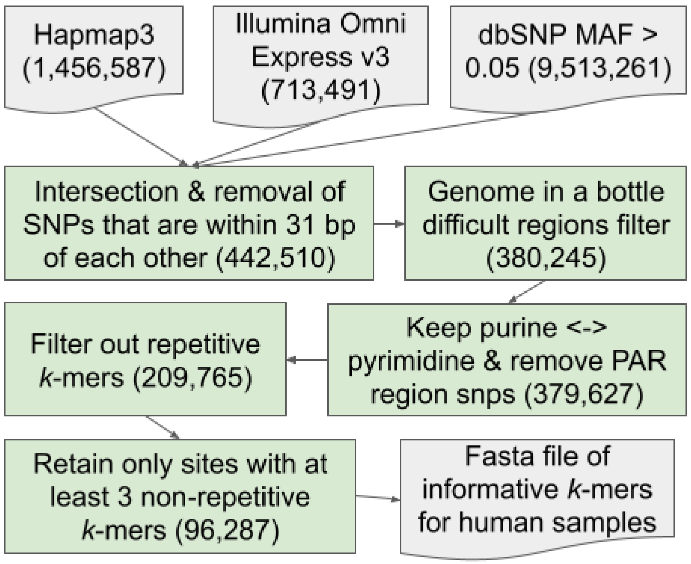


Figure 1. The selection method for informative human polymorphic sites used in *k*-mer counting based sample swap detection.

#### Generation PCA rotation matrices for human samples

In addition to variant sites sequences themselves, ntsm can optionally use population derived PCA rotational matrices which can help speed up comparisons of a large number of samples. We provide a python script that utilizes pandas [23] and scikit-learn [24] for those who wish to generate their own rotational matrices from a multiVCF file.

Using the multiVCF file from the 1000 Genome project [25], we generated a matrix of samples to our selected variant sites above (0 for homozygous A/T allele, 0.5 for heterozygous alleles, and 1 for homozygous C/G alleles). This is followed by normalizing the matrix by the standard deviation for each site and we keep the normalization vector in a file. This sample to variant matrix undergoes decomposition into principal components (Figure 2), though instead of being concerned with the principal component values themselves, we are primarily interested in keeping the rotational matrices for a number of the most significant components, and using them to project the new sequences onto.

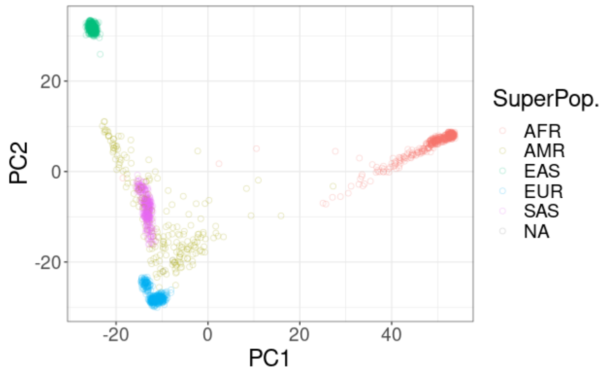


Figure 2. A population PCA of the first 2 components using the variant human sites selected and the 1000 Genomes Project multiVCF dataset. Each color represents a different labeled superpopulation group.

### Implementation Details

#### Variant *k*-mer Counting

Paired variant sequences (1 file for C/G allele and 1 file for A/T variants) are stored in fasta files before being loaded. These alleles are then broken into *k*-mers and hashed using an invertible hash function into a hash table [26]. A sliding *k*-mer window for each allele is used, to provide redundancy to compensate for sequencing errors. Input sequences in fastq format are then read, broken in *k*-mers and also hashed [27], then subsequently checked for existence in the hash table. If they exist, then the occurrence count for that *k*-mer increments by one.

Optionally, the total number of *k*-mers that match a site can be used as criteria for early termination to save computational time, as only fairly low coverage is needed to perform accurate classification. For each allele, only the highest count of the sequence is outputted in the results counts file. The counts are a simple tab-separated (TSV) file with 3 columns for the site identifier, allele A/T count and allele C/G count. Multiple files and threads can be used on the same instance of the counting tool for speed, or run separately on different files and later merged.

#### Sequencing error rate estimation

Error rate can inform the user of the viability of the dataset and inform the user as to why downstream applications may be performing poorly. In our case, it may help explain why a sample swap signal is weak.

To estimate the error rate our tool records the total count of all *k*-mers seen *t* and the total number of *k*-mers matching to our set of *k*-mers *m*. To relate these values to each other, we need an expected number of *k*-mers *n* assuming no error. Assuming that the data is randomly sampled from the genome, we can find the expected value of *n* given the diploid genome size *g* and the total number of distinct *k*-mers within our set of *k*-mer *d* with the following formula:

$$\hat{n} = \frac{td}{g}$$

Using *n*, we can use maximum likelihood estimation [28] (MLE) to derive the estimate the expected similarity *p*:

$$\mathcal{L}(p) = \left(p^k\right)^m \left(1 - p^k\right)^{n-m} = p^{km} \left(1 - p^k\right)^{n-m}$$

Working in log space will make our MLE derivation easier,

$$\log \mathcal{L}(p) = m \cdot \log\left(p^k\right) + (nm) \cdot \log\left(1 - p^k\right)$$

Thus,

$$\frac{\partial \log \mathcal{L}(p)}{\partial p} = \frac{km}{p} - \frac{n-m}{1-p^k} \cdot kp^{k-1} = \frac{k \cdot \left[m(1-p^k) - (n-m)p^k\right]}{p\left(1-p^k\right)}$$

The maximum likelihood estimate of *p* is obtained when $\partial \log \mathcal{L} / \partial p = 0$. Thus,

$$0 = \frac{k \cdot \left[m - mp^k - np^k + mp^k\right]}{p\left(1-p^k\right)}$$

$$0 = m - np^k$$

Finally, similarity is formulated as:

$$\hat{p} = \left(\frac{m}{n}\right)^{1/k}$$

Error rate is merely the inverse of similarity,

$$ErrorRate = 1 - \left(\frac{m}{n}\right)^{1/k}$$

We note that this formulation largely holds true for mismatch and small indel error types, however when large indels are introduced this formulation can become less accurate depending on how one defines the ground truth alignment used to calculate the sequence error rate.

#### Similarity score for detecting sample swaps

Rather than computing genotypes for each site, our method directly uses the counts of each allele derived from k-mer counts. Our score is derived using likelihood ratio test as a basis with 2 models assuming that either the samples are independent or if they are the same sample. To minimize the effect of missing data due to low coverage, in each pairwise analysis we remove sites with missing counts for both alleles. The score is further modulated to be extra conservative, lowering the confidence of our result when coverage of the dataset is low.

To start, we need the likelihood of one sample. Suppose there are *N* sites. For a sample at site *i*, we observe $x_{ia}$ count of allele *a*, where $a \in \{1, 2\}$ for the two alleles at each site *i*. Let $p_{ia}$ be the probability of observing allele *a* at site *i*. Then the probability of the data **x** is:

$$\mathcal{L}(\mathbf{p}) = P(\mathbf{x}|\mathbf{p}) = \prod_{i=1}^{N}\prod_{a=1}^{2} p_{ia}^{x_{ia}}$$

The log-likelihood is

$$\log \mathcal{L}(p) = \sum_{i}\sum_{a} x_{ia} \log p_{ia}$$

The max-likelihood (ML) estimate of $p_{ia}$ is

$$\hat{p}_{ia} = \frac{x_{ia}}{x_{i1} + x_{i2}}$$

Next, for a log-likelihood ratio test, we need to compare two models:

Model 1: two samples are independent
Let $L^{(1)}$ be the likelihood of sample 1 and so it is with sample 2. The total likelihood is $L^{(1)} \cdot L^{(2)}$.

Model 2: two samples are the same
In this case, we can merge all counts. Let

$$x_{ia}^{(*)} = x_{ia}^{(1)} + x_{ia}^{(2)}$$

Then the probability of the two sample is

$$P^{(*)} = \prod_{i=1}^{N}\prod_{a=1}^{2} \left[p_{ia}^{(*)}\right]^{x_{ia}^{(*)}}$$

The ML estimate of $p_{ia}^{(*)}$ is

$$\hat{p}_{ia}^{(*)} = \frac{x_{ia}^{(*)}}{x_{i1}^{(*)} + x_{i2}^{(*)}}$$

and the log-likelihood is

$$\log \mathcal{L}^{(*)}(p) = \sum_{i}\sum_{a} x_{ia}^{(*)} \log p_{ia}^{(*)}$$

Using the two models proposed previously we can input the results into the Log-likelihood ratio test [29]. The log likelihood ratios can be used to compute a robust score metric which can determine if two samples are of the same origin. The log-likelihood formulation is as follows:

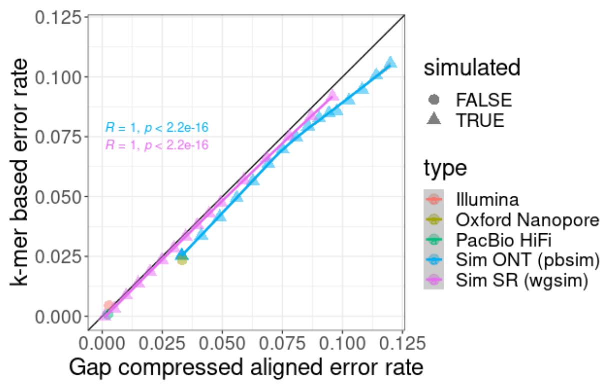


Figure 3. *k*-mer based error rate calculation vs Gap compressed align error rate ground truth on real and simulated CHM13 data onto the CHM13 T2T reference genome.

We found that our estimates closely match the expected error rate (Fig. 3), though both the real and simulated ONT data was slightly underestimated on average, but not to a degree that makes the estimate unreliable. It is expected that error caused by indels, especially if long segments of these are present, would produce a lower calculated error rate as the formulation (see methods) for our error rate calculator assumes only mismatches can occur. That said, we expected the formulation's the short indels to contribute to the error calculation  in a way very similar to mismatches which is reflected here. We note that for our method an estimated diploid genome size is needed, and for these tests the default value used was 6.2 Gb. This value of course will differ if a genome with a different size genome is used.

#### PCA-based heuristic investigation

To evaluate the efficacy of our PCA-based method to reduce the number of pairwise comparisons that we perform, we use data from the HPRC (Supplementary Table S1), and run them with ntsm while providing a rotation matrix with normalization values. The properties of the data varied wildly between coverage and error rate providing comprehensive expected performance of our heuristics given different data types.

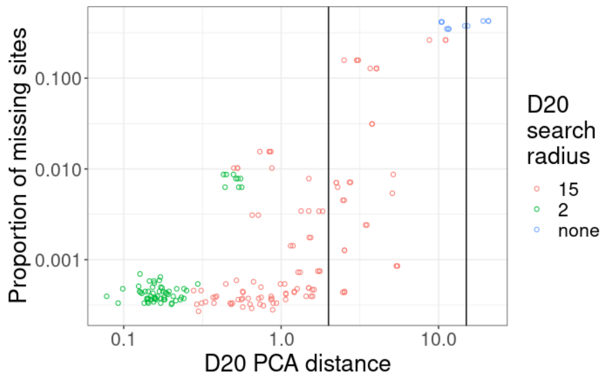


Figure 4. The PCA distance (top 20 PCs) between matching sample pairs vs the proportion of missing sites found in the dataset with the most missing sites. The colors (D20 search radius) correspond to the radius chosen by the algorithm for each sample pair based on the error rate and number of missing sites The vertical lines denote the radius thresholds of 2 and 15.

The number of missing sites greatly affects the performance of our heuristic, generally requiring a larger radius to search as the number of missing sites increase (Figure 4). We can measure the number of missing sites to apply thresholds prior to applying a search radius. The other variable determining the performance of our heuristic is the correctness of our genotype calls on our data. This is influenced by both the coverage of the data and error rate of the data. We also use our error rate estimates to determine the radius to search. The coverage is largely a function of the missing sites and found it to be a better metric overall. Because of this, high coverage data generally requires a much smaller search radius (Supp. Figure S2).

When the *k*-mer coverage smaller dataset of a pair of matching samples >20x, we find that >57% of the samples are set to use a radius of 2, while at a coverage >30x, we find that >93% of the samples require a search radius of 2 (the remaining datasets at search radius of 15 being higher error nanopore data). At a search radius of 15 the expected number of candidates is <25% of the total number of samples, and at 2 the expected number of candidates to search drops to <5% of possible elements on average (Figure 5).

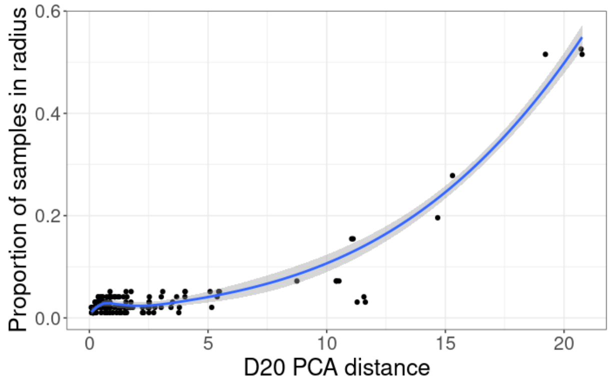


Figure 5. The PCA distance between each matching sample pair and the average proportion of samples of that are determined as candidates for at that radius.

### Comparisons to Somalier

To show the performance of ntsm, we sequence data from the Human Pangenome Reference Consortium (HPRC) [17] and a multiVCF file from the 1000 Genomes Project [25] featuring 3202 samples. We compare 39 samples with whole genome data from the HPRC which include sequencing data from Illumina, Pacbio HiFi, Hi-C, 10x Chromium, Strand-seq, and Oxford Nanopore platforms (Supplementary Table S1). As we expected Oxford Nanopore to be the most difficult datatype to analyze, we chose only samples that had complementary data of this type in the analysis.

#### Sensitivity and Specificity of Sample Swaps

Unlike ntsm, Somalier does not provide an automated means of determining what should be considered similar enough to consider it the same sample. However, it does provide relatedness metrics that can easily be used to threshold samples showing high degrees of relatedness and thus similarity. To determine a threshold, we ran the full coverage of each dataset for Somalier and manually picked a threshold that provided perfect separations between samples with the same origins from those with different origins (Figure 6). We determined that a relatedness value of 0.667 seemed to be a good threshold while still maintaining a high sensitivity. As mentioned in the methods section, ntsm uses a log-likelihood based score to separate samples with same origins from those with different origins, where the default value for this threshold is 0.5.

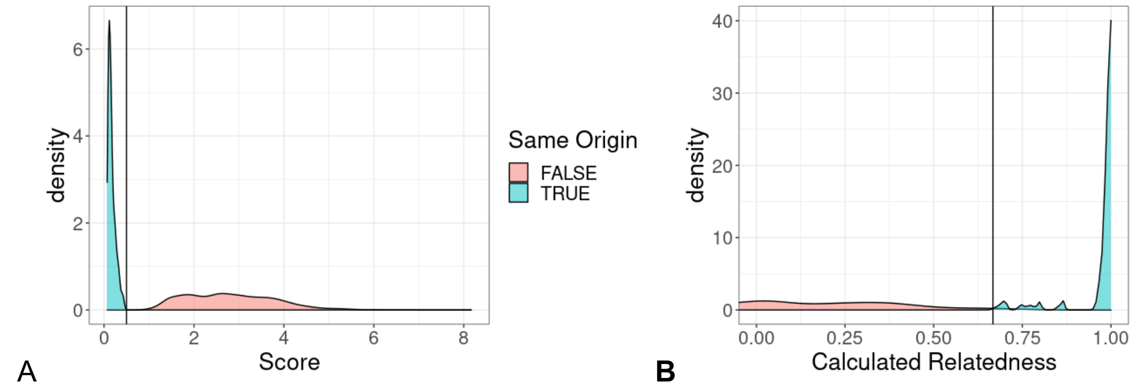


Figure 6. Distributions of metrics (score for ntsm (B) and relatedness for Somalier (A)) used for determining if two samples are similar. Horizontal lines are the thresholds used to determine the same sample status in later parts of the comparison (Relatedness > 0.67 for Somalier and Score < 0.5 for ntsm, respectively).

Next we randomly subsampled each dataset at different fold coverages (from 0.5x to 20x) and proceeded to see how each tool performed on this lower coverage data. We found that both tools are capable of detecting whether or samples have the same sample of origin at coverages higher than 5x, however Somalier struggled when attempting to match samples at coverage lower than 5x, producing a much higher number of false positive and negative pairs in the output (Figure 7). At sub-1x coverage, even ntsm struggled with detecting samples with the same origin, though not to the degree that Somalier struggled. In general, ntsm benefits from higher coverage as well, generating higher scores (i.e. more confident results) for unmatched samples (Supplementary Figure S1.).

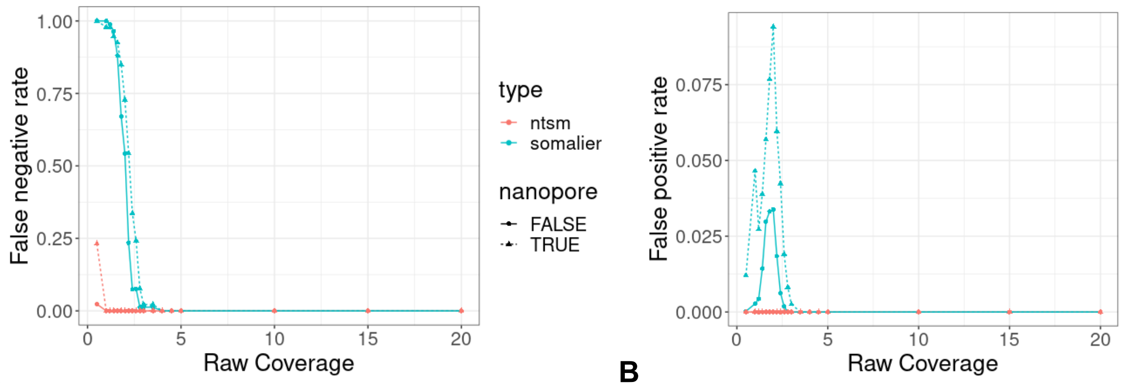


Figure 7. Effects of dataset coverage on classification performance on Somalier and ntsm. A. The false negative rate (FNR) of Somalier and ntsm at varying raw dataset coverage. B. The false positive rate (FPR) of Somalier and ntsm at varying raw dataset coverage. Nanopore combined with other datasets is shown separately due to the high error rate of the former.

#### Memory and time comparisons

##### *k*-mer counting (ntsm) vs alignment (somalier)

One of the primary benefits of ntsm is bypassing the alignment requirement that other tools require. However, the alternative we must perform is *k*-mer counting, which, though is fairly resource frugal, is not free. To determine the relative resource cost counting takes in comparison to alignment, we took equal coverage subsamples (2x) of our datasets (Supplementary Table S1) and ran them with ntsm and each alignment tool we used to generate alignments needed for Somalier.

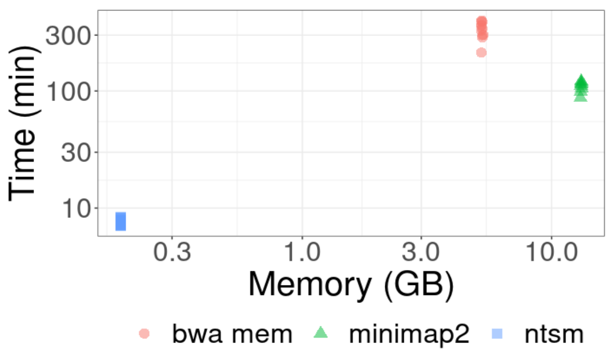


Figure 8. Peak memory and wallclock time comparisons of *k*-mer counting vs alignment using single thread

We then measured the time and memory needed for each tool used (Figure 8). We found that ntsm ran at an average of ~8 minutes, orders of magnitude less than bwa mem and minimap2 at ~1.9 hours and 5.9 hours respectively. Memory usage is low because we are only counting a very small specific subset of *k*-mers. Note that we did not include sorting or indexing time in this analysis as we hoped to illustrate that even without this in our comparison *k*-mer counting was still much less resource intensive. Also sorting can partially be run in parallel with alignments as reads are streamed.

##### Sample comparison process

As mentioned in the method section, comparison of all samples with each other is naively an all-to-all operation and thus a quadratically scaling operation. For most studies this may still be a trivial concern but as larger and larger studies are considered, this can become an increasingly expensive consideration. To observe the computational expense of this process we took subsamples of the 1000 Genomes VCF files and measured time and memory used to compute relatedness between the samples.

As expected we observed that pairwise comparison time scaled quadratically (Figure 9), even in Somalier, albeit with each individual comparison being orders of magnitude faster than our method. Somalier, utilizing a bitvector based comparison method is much more optimized for speed than our count based method, however we show that our PCA-based screening method is still competitive. We observe here that our PCA-based screen approach actually may scale less than quadratically though it requires high coverage and low error rate data to perform this way reliably. Memory usage is largely linear relative to the data (Supp. Figure S3) as expected and there is no additional memory overhead using our PCA-based method.

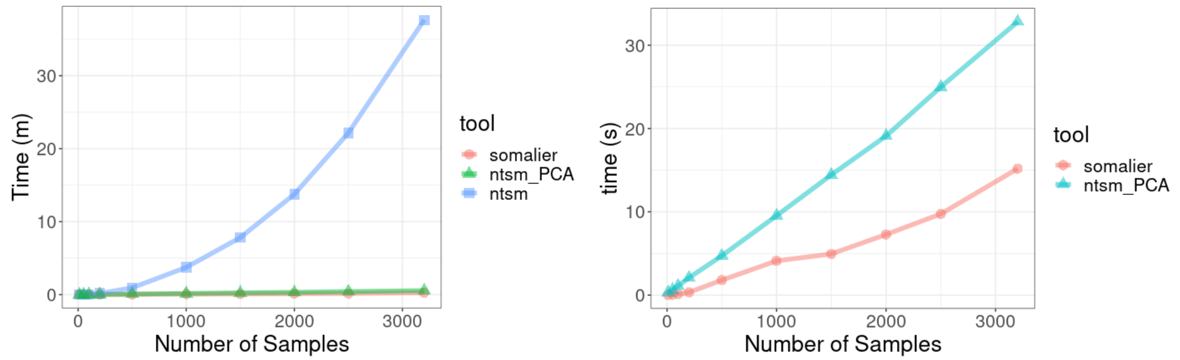


Figure 9. Wall clock time comparisons of somalier vs ntsm with and without PCA-based screening. A multiVCF file for 3202 samples from the 1000 Genome project was used. Somalier was capable of only using 1 thread while ntsm used 16 threads.

#### *k*-mer based relatedness calculation

To test our relatedness estimation methods we took Pacbio HiFi datasets of the parent (HG003, HG004) child (HG002) trio. Both ntsm and Somalier correctly computed the relatedness we expected; that is, parents remained unrelated (0%), while child samples showed 50% relatedness to its parents and with 100% relatedness to a technical replicate to itself (Figure 10)

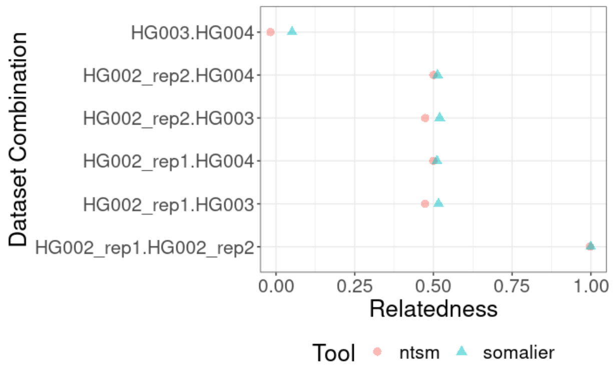


Figure 10. Comparison of Somalier and ntsm computed relatedness  of parent child trio (HG002, HG003, HG004) from Pacbio HiFi sequencing reads. Two independent sequencing runs of HG002 are used here.

Though the similarity values largely agree between ntsm and Somalier, there are minor differences between our calculations. These differences likely primarily stem from the fact that Somailer and ntsm use different variant sites and that we use *k*-mer counts to create genotyping calls and omit missing sites from the relatedness calculation. The differences in our tool are likely due to the number sites used by default by somalier (total of 17,766 sites), and the sites we selected (see methods, total of 96,287 sites). In addition, some of the differences in our relatedness calculation are a consequence of omitting missing sites, which Somalier is unable to perform as it computes similarity using bit vectors of a uniform length.

To gain a more comprehensive view on the accuracy of our relatedness calculations we checked the quality of our relatedness estimates using samples shown to be related in 1000 Genomes cohorts in addition supplemented with samples with at least 20x coverage sequence coverage of various sequencing data types (Supp. Table 1). We found that overall ntsm produces a relatedness metric closer and more tightly grouped to the expected value based on the pedigree (Figure 11). We note however this trend does not hold true for relatedness estimates involving a nanopore dataset, showing that ntsm calculates relatedness conservatively when it comes to data originating from the same sample.

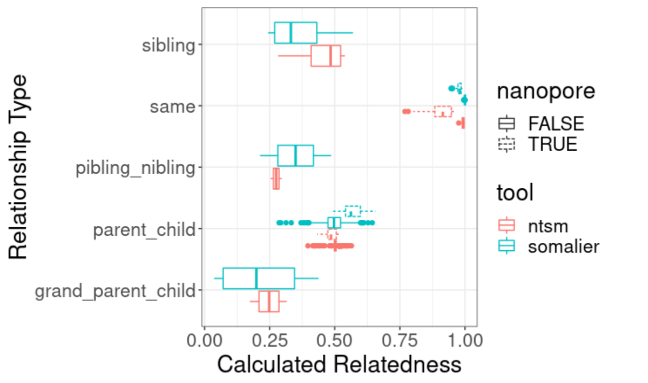


Figure 11. Comparison of known related samples in 1000 Genomes cohort, supplemented with samples with ~20x coverage sequence coverage of various sequencing data types (Supp. Table 1). Relatedness calculations involving a Nanopore dataset are shown separately.

### References


1. Chu J, Sadeghi S, Raymond A, Jackman SD, Nip KM, Mar R, et al.. BioBloom tools: fast, accurate and memory-efficient host species sequence screening using bloom filters. *Bioinformatics*. 30:3402–42014;

2. Cornet L, Baurain D. Contamination detection in genomic data: more is not enough. *Genome Biol*. BioMed Central; 23:1–152022;

3. Ondov BD, Treangen TJ, Melsted P, Mallonee AB, Bergman NH, Koren S, et al.. Mash: fast genome and metagenome distance estimation using MinHash. *Genome Biol*. 17:1322016;

4. Manichaikul A, Mychaleckyj JC, Rich SS, Daly K, Sale M, Chen WM. Robust relationship inference in genome-wide association studies. *Bioinformatics*. Bioinformatics; 2010; doi: 10.1093/bioinformatics/btq559.

5. Bergmann EA, Chen B-J, Arora K, Vacic V, Zody MC. Conpair: concordance and contamination estimator for matched tumor–normal pairs. *Bioinformatics*. Oxford Academic; 32:3196–82016;

6. Wang PPS, Parker WT, Branford S, Schreiber AW. BAM-matcher: a tool for rapid NGS sample matching. *Bioinformatics*. Oxford Academic; 32:2699–7012016;

7. Schröder J, Corbin V, Papenfuss AT. HYSYS: have you swapped your samples? *Bioinformatics*. Oxford Academic; 33:596–82016;

8. Lee S, Lee S, Ouellette S, Park W-Y, Lee EA, Park PJ. NGSCheckMate: software for validating sample identity in next-generation sequencing studies within and across data types. *Nucleic Acids Res*. Oxford Academic; 45:e103–e1032017;

9. Pedersen BS, Quinlan AR. Who's Who? Detecting and Resolving Sample Anomalies in Human DNA Sequencing Studies with Peddy. *Am J Hum Genet*. Elsevier; 100:406–132017;

10. Javed N, Farjoun Y, Fennell TJ, Epstein CB, Bernstein BE, Shoresh N. Detecting sample swaps in diverse NGS data types using linkage disequilibrium. *Nat Commun*. Nature Publishing Group; 11:1–82020;

11. Pedersen BS, Bhetariya PJ, Brown J, Kravitz SN, Marth G, Jensen RL, et al.. Somalier: rapid relatedness estimation for cancer and germline studies using efficient genome sketches. *Genome Med*. 12:622020;

12. Bennett S. Solexa Ltd. *Pharmacogenomics*. Pharmacogenomics; 2004; doi: 10.1517/14622416.5.4.433.

13. Branton D, Deamer DW, Marziali A, Bayley H, Benner SA, Butler T, et al.. The potential and challenges of nanopore sequencing. *Nat Biotechnol*. NIH Public Access; 26:11462008;

14. Belton J-M, McCord RP, Gibcus J, Naumova N, Zhan Y, Dekker J. Hi-C: A comprehensive technique to capture the conformation of genomes. *Methods*. NIH Public Access; 2012; doi: 10.1016/j.ymeth.2012.05.001.

15. Zhang M, Zhang Y, Scheuring CF, Wu C-C, Dong JJ, Zhang H-B. Preparation of megabase-sized DNA from a variety of organisms using the nuclei method for advanced genomics research. *Nat Protoc*. Nature Publishing Group; 7:467–782012;

16. Rhie A, McCarthy SA, Fedrigo O, Damas J, Formenti G, Koren S, et al.. Towards complete and error-free genome assemblies of all vertebrate species. *Nature*. 592:737–462021;

17. Liao W-W, Asri M, Ebler J, Doerr D, Haukness M, Hickey G, et al.. A draft human pangenome reference. *Nature*. 617:312–242023;

18. . The International HapMap Project. *Nature*. Nature Publishing Group; 426:789–962003;

19. : [No title]. https://www.illumina.com/Documents/products/datasheets/datasheet_gwas_roadmap.pdf Accessed 2023 Oct 27.

20. Smigielski EM. dbSNP: a database of single nucleotide polymorphisms. Nucleic Acids Research.

21. Zook JM, Catoe D, McDaniel J, Vang L, Spies N, Sidow A, et al.. Extensive sequencing of seven human genomes to characterize benchmark reference materials. Sci Data. p. 160025.

22. Li. Aligning new-sequencing reads by BWA. *Broad Institute*.

23. The pandas development team. pandas-dev/pandas: Pandas. Zenodo;

24. Garreta R, Moncecchi G. Learning Scikit-Learn: Machine Learning in Python. Packt Pub Limited;

25. 1000 Genomes Project Consortium, Auton A, Brooks LD, Durbin RM, Garrison EP, Kang HM, et al.. A global reference for human genetic variation. *Nature*. 526:68–742015;

26. : Website. https://github.com/Tessil/robin-map)

27. : Integer Hash Function. http://web.archive.org/web/20071223173210/http://www.concentric.net/~Ttwang/tech/inthash.htm Accessed 2023 Sep 8.

28. Fisher RA. On the mathematical foundations of theoretical statistics. *Philos Trans R Soc Lond*. The Royal Society; 222:309–681922;

29. Wilks SS. The large-sample distribution of the likelihood ratio for testing composite hypotheses. *Ann Math Stat*. Institute of Mathematical Statistics; 9:60–21938;

30. Patterson N, Price AL, Reich D. Population structure and eigenanalysis. *PLoS Genet*. 2:e1902006;

31. Bentley JL. Divide and Conquer Algorithms for Closest Point Problems in Multidimensional Space.

32. : GitHub - jlblancoc/nanoflann: nanoflann: a C++11 header-only library for Nearest Neighbor (NN) search with KD-trees. GitHub. https://github.com/jlblancoc/nanoflann Accessed 2023 Oct 30.

33. Nurk S, Koren S, Rhie A, Rautiainen M, Bzikadze AV, Mikheenko A, et al.. The complete sequence of a human genome. *Science*. 376:44–532022;

34. Danecek P, Bonfield JK, Liddle J, Marshall J, Ohan V, Pollard MO, et al.. Twelve years of SAMtools and BCFtools. *Gigascience*. Oxford Academic; 10:giab0082021;

35. Ono Y, Asai K, Hamada M. PBSIM2: a simulator for long-read sequencers with a novel generative model of quality scores. *Bioinformatics*. Oxford Academic; 37:589–952020;

36. Li H: On the definition of sequence identity. https://lh3.github.io/2018/11/25/on-the-definition-of-sequence-identity Accessed 2023 Oct 27.

37. Albert TJ, Molla MN, Muzny DM, Nazareth L, Wheeler D, Song X, et al.. Direct selection of human genomic loci by microarray hybridization. *Nat Methods*. Nat Methods; 2007; doi: 10.1038/nmeth1111.

38. Wang Z, Gerstein M, Snyder M. RNA-Seq: a revolutionary tool for transcriptomics. *Nat Rev Genet*. 10:57–632009;

39. Johnson DS, Mortazavi A, Myers RM, Wold B. Genome-wide mapping of in vivo protein-DNA interactions. *Science*. Science; 2007; doi: 10.1126/science.1141319.

40. Brown CT, Howe A, Zhang Q, Pyrkosz AB, Brom TH. A reference-free algorithm for computational normalization of shotgun sequencing data. arXiv; 2012; doi: 10.48550/ARXIV.1203.4802.

41. Gu J, Dai J, Lu H, Zhao H. Comprehensive Analysis of Ubiquitously Expressed Genes in Humans from A Data-driven Perspective. *Genomics Proteomics Bioinformatics*. 21:164–762023;

### Supplement

#### Read sequence dataset information

| Sample ID | Hi-c | Nanopore | Strand seq | Illumina | PacBio HiFi | 10X |
|---|---|---|---|---|---|---|
| HG002 | 84.13 | 82.88 | 6.77 | 30.51 | 39.64 | - |
| HG00438 | 82.2 | 39.08 | - | 30.33 | 29.19 | - |
| HG005 | - | 103.13 | - | 47.02 | 44.01 | - |
| HG00621 | 83.7 | 30.06 | - | 29.36 | - | - |
| HG00673 | 86.09 | 32.83 | - | 28.7 | - | - |
| HG00733 | 57.12 | 62.31 | - | 30.89 | 32.19 | - |
| HG00735 | 72.78 | 29.35 | - | 30.04 | - | - |
| HG00741 | 68.35 | 41.96 | - | 31.1 | - | - |
| HG01071 | 68.6 | 37.71 | - | 31 | - | - |
| HG01106 | 64.23 | 38.8 | - | 31.17 | - | - |
| HG01109 | 19.01 | 58.78 | - | 30.56 | 30.18 | 59.76 |
| HG01123 | 52.97 | 24.47 | 1.65 | - | 37.15 | - |
| HG01175 | 68.64 | 42.14 | - | 29.2 | 34.83 | - |
| HG01243 | 26.97 | 50.57 | - | 29.71 | 33.16 | 53.05 |
| HG01258 | 62.85 | 19.76 | - | 31.49 | - | - |
| HG01358 | 53.63 | 14.67 | 2.5 | 31.5 | - | - |
| HG01361 | 46.74 | 30.67 | - | 32.64 | - | - |
| HG01891 | 41.53 | 34.35 | 1.57 | 30.6 | - | - |
| HG01928 | 68.75 | 22.85 | - | 32.66 | - | - |
| HG01952 | 61.03 | 29.18 | - | 31.26 | - | - |
| HG01978 | 85.89 | 36.74 | - | 33.56 | - | - |
| HG02055 | 28.8 | 52.38 | - | 30.65 | 37.39 | 58.32 |
| HG02080 | 28.78 | 47.63 | - | 31.92 | 32.89 | 45.07 |
| HG02148 | 63.77 | 17.39 | - | 29.09 | - | - |
| HG02257 | 50.71 | 18.43 | 3.98 | 31.92 | - | - |
| HG02486 | 38.47 | 27.02 | 1.96 | - | - | - |
| HG02559 | 48.27 | 22.59 | - | - | - | - |
| HG02572 | 115.82 | 14.59 | - | 30.37 | - | - |
| HG02622 | 52.18 | 13.56 | - | 33.3 | - | - |
| HG02630 | 54.73 | 18.36 | - | 33.31 | - | - |
| HG02717 | 72.3 | 25.38 | - | 32.68 | - | - |
| HG02886 | 56.86 | 30.7 | - | 34.2 | - | - |
| HG03098 | 23.01 | 44.57 | - | 30.66 | 33.55 | 57.83 |
| HG03453 | 60.4 | 10.29 | - | 29.93 | - | - |
| HG03471 | 58.84 | 27.46 | - | - | - | - |
| HG03492 | 30.48 | 40.36 | - | 29.19 | 31.61 | 56.05 |
| HG03516 | 49.44 | 40.25 | - | 28.65 | - | - |
| HG03540 | 52.88 | 27.05 | - | 33.81 | - | - |
| HG03579 | 69.9 | 14.8 | - | 29.06 | - | - |

Supplementary Table S1. HPRC sequencing datasets used in analysis. Each column after the first represents the aligned coverage (either using minimap2 or bwa mem) of each data type available for analysis.

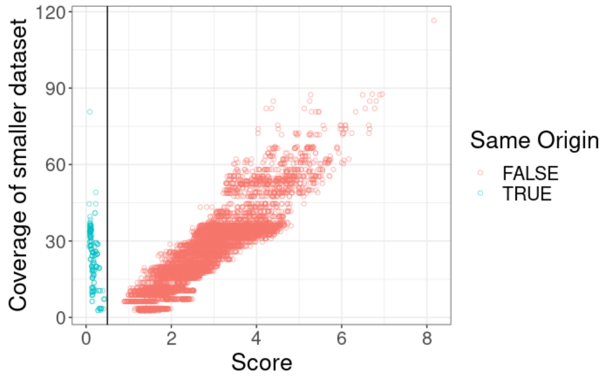


Supplementary Figure S1. Comparison of score and *k*-mer coverage on samples from the same origin and not. Confidence in distinguishing samples increases as coverage increases, reflected in high scores the more distant they are.

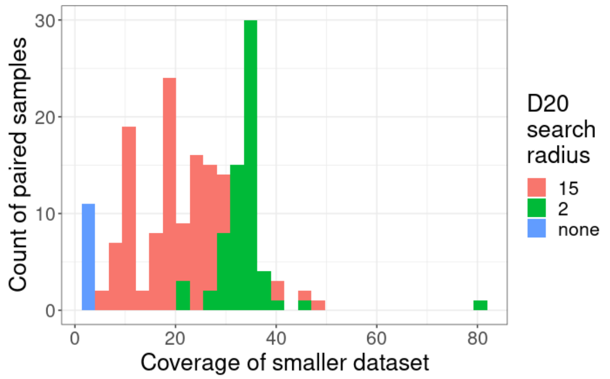


Supplementary Figure S2. The *k*-mer coverage of the smaller sample in a matching sample pair and the number of sample pairs that use a specific search radius.

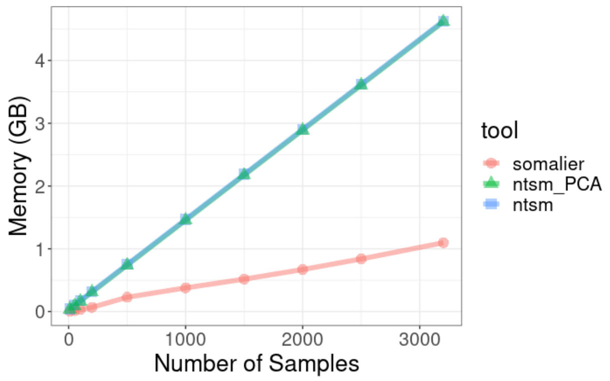

Supplementary Figure S3. The memory of somalier vs ntsm using with and without PCA-based prescreening, processing similarity between 3202 samples from the 1000 Genome project.
